# Vitamin D3 regulates estrogen’s action and affects mammary epithelial organization in 3D cultures

**DOI:** 10.1101/439307

**Authors:** Nafis Hasan, Carlos Sonnenschein, Ana M. Soto

## Abstract

Vitamin D3 (vitD3) and its active metabolite, calcitriol (1,25-(OH)_2_D_3_), affect multiple tissue types by interacting with the vitamin D receptor (VDR). Although vitD3 deficiency has been correlated with increased incidence of breast cancer and less favorable outcomes across ethnic groups and latitudes, randomized human clinical trials have yet to provide conclusive evidence on the efficacy of vitD3 in treating and/or preventing breast cancer. When considering that carcinogenesis is “development gone awry”, it becomes imperative to understand the role of vitD3 during breast development. Mammary gland development in VDR KO mice is altered by increased ductal elongation and lateral branching during puberty, precocious and increased alveologenesis at pregnancy and delayed post-lactational involution. These developmental processes are largely influenced by mammotropic hormones, i.e., ductal elongation by estrogen, branching by progesterone and alveologenesis by prolactin. However, research on vitD3’s effects on mammary gland morphogenesis focused on cell proliferation and apoptosis in 2D culture models and utilized supra-physiological doses of vitD3, conditions that spare the microenvironment in which morphogenesis takes place. Here, using two 3D culture models, we investigated the role of vitD3 in mammary epithelial morphogenesis. We found that vitD3 interferes with estrogen’s actions on T47D human breast cancer cells in 3D differently at different doses, and recapitulates what is observed *in vivo*. Also, vitD3 can act autonomously and affect the organization of MCF10A cells in 3D collagen matrix by influencing collagen fiber organization. Thus, we uncovered how vitD3 modulates mammary tissue organization independent of its already known effects on cell proliferation.

## Introduction

Breast cancer remains a major cause of mortality among women worldwide. Epidemiological studies have shown that key stages during breast development are particularly susceptible to the effects of carcinogens. For instance, women aged 10-19 years who were exposed to atomic bomb radiation in Hiroshima in World War II showed an excess of breast cancer cases at the age of prevalence compared to similarly exposed women aged 35 years and older (1). Likewise, women exposed to diethylstilbestrol during fetal life have a higher risk of breast cancer compared to unexposed women (2), and women exposed to DDT in the womb showed a four-fold higher risk of breast cancer in adulthood (3). This phenomenon has also been observed in rodents; namely, rats exposed to NMU around puberty have a 100% incidence of tumors, but the incidence rate falls to just 10% when exposed after 90 days of age (4). Rodents exposed *in utero* to low doses of BPA have also shown a higher incidence of mammary gland tumors in adult life (5,6). These “windows of susceptibility” coincide with key milestones of organogenesis and/or tissue remodeling, buttressing the notion that carcinogenesis is “development gone awry.”(7)

Vitamin D3 (VitD3), and its active metabolite calcitriol (1,25-(OH)_2_D_3_), has been primarily studied in the context of normal and diseased bone development (8). However, research over the past few decades have shown that vitD3 can affect multiple tissue types, including the mammary gland (8). For instance, epidemiological data from human populations across different ethnicities and latitudes show a correlation between vitD3 deficiency and risk of breast cancer incidence, and between vitD3 deficiency and worse outcomes for breast cancer patients (8–10). Animal models show that calcitriol deficiency promotes tumor growth in mice (11), and vitamin D receptor (VDR) KO animals have a higher mortality rate when they are crossed into a genetically induced breast cancer model background (12). Moreover, dietary supplementation of vitD3 inhibited tumor growth in xenograft models of breast and prostate cancers in mice (13). Although vitD3 supplementation may help with patient outcomes, and post-menopausal women may benefit from a lower breast cancer risk (14), there is yet no conclusive evidence from randomized clinical trials that validates the efficiency of vitD3 as a therapeutic or preventive option for breast cancer.

Mammary glands from VDR KO mice exhibit a florid developmental phenotype, as exemplified by increased number of terminal end buds (TEBs), increased ductal extension and lateral branching at puberty (15), precocious alveologenesis during pregnancy, and delayed post-lactational involution (16). These developmental processes are largely influenced by the so-called mammotropic hormones, i.e., Estradiol (E2), Progesterone (Prg) and Prolactin (Prl). More specifically, E2 induces ductal elongation, Prg increases lateral branching and Prl stimulates alveologenesis (17). The VDR KO phenotype in the mammary gland of rodents indicates that vitD3 interacts with these hormones during mammary epithelial morphogenesis. However, no study has yet examined the interactions between these mammotropic hormones and vitD3 at these developmental stages, or explored the role of vitD3 in mammary epithelial morphogenesis.

Studies on vitD3’s action in the mammary gland have largely focused on vitD3’s effects on cell proliferation and apoptosis (8). These studies have mostly utilized 2D cell culture models; they do not represent the 3D environment required for morphogenesis. Additionally, most studies have used a calcitriol dose of 100 nM. This dose was chosen based on the deficiency cut-off level for vitD3 in human populations, which is determined by measuring the serum levels of calcidiol, the precursor to calcitriol (18). Therefore, utilizing such a dose assumes a hundred percent conversion rate of calcidiol to calcitriol locally at specific tissues, and does not take into account that calcitriol may elicit a non-monotonic dose response comparable to that observed with other steroid hormones.

In this study, we have utilized two different 3D cell cultures models to investigate vitD3’s role in mammary gland morphogenesis. We observed that calcitriol constrains E2’s actions on the organization of mammary epithelial cells, and that this effect is independent of its already well-characterized role in cell proliferation. We also noticed that calcitriol has autonomous effects on mammary ductal organization and describe a novel effect of calcitriol on the organization of collagen fibers. Finally, we confirm that calcitriol elicited a non-monotonic dose response in mammary epithelial cells, as can be expected from a steroid hormone, and therefore confirmed that vitD3 acts as a steroid hormone.

## Materials & Methods

### Reagents

Hydrocortisone, cholera toxin, insulin, Dulbecco’s modified Eagle’s medium (DMEM), Phosphate Buffered Saline (PBS, 10x), and calcitriol were purchased from Sigma-Aldrich (St. Louis, MO). DMEM/F12, L-Glutamine and Trypsin were purchased from Life Technologies (Carlsbad, CA). 17β-estradiol (E2) was purchased from Calbiochem. Fetal bovine and equine sera were purchased from Thermo Fisher (Waltham, MA). Epidermal Growth Factor (EGF) and rat tail type I collagen were purchased from Corning (Tewksbury, MA). E2 and calcitriol were resuspended in ethanol to make 10^−3^ and 10^−4^ M stocks, respectively.

### Cell maintenance

Human breast epithelial T47D cells used in this study were cloned from a population originally obtained from Dr. G. Green (U. Chicago). The cells used in these experiments were tested for estrogen sensitivity before they were used (19). These cells were grown in the DMEM containing 5% FBS (propagation medium). When looking at effects of hormones on these cells using 2D culture experiments, we used DMEM/F12 without phenol red with 5% charcoal-dextran stripped FBS (CD-FBS) and 10^6^ U/ml penicillin; when performing 3D culture experiments with the same purpose, we used a mixture of 75% DMEM, 25% Ham’s F12 without phenol red, 7.5% CD-FBS and 2 mM L-glutamine. Human breast epithelial MCF10A cells were purchased from American Type Culture Collection (Manassas, VA) and maintained in DMEM/F12 with phenol red, 5% equine serum, 20 ng/ml epidermal growth factor (EGF), 0.5 μg/ml hydrocortisone, 0.1 μg/ml cholera toxin, and 10 μg/ml insulin. Experiments performed using MCF10A cells used the same medium. In all cases, cells were incubated at 37°C in 6% CO_2_ and 100% humidity.

### Dose response curves to calcitriol

Dose-response curves to calcitriol in 2D culture were performed in 24-well plates. Cells were seeded at a density of 35,000/well in media and allowed to attach. For MCF10A cells, seeding media was changed to media containing different doses of calcitriol after 24 hours of seeding. For T47D cells, propagation media was removed 48 hours after seeding, and substituted with CD-FBS medium containing different doses of calcitriol with or without E2. After 6 days, cell numbers were determined using the SRB assay (20).

### 3D cultures

Collagen type I gels were formulated at a final concentration of 1 mg/mL as described previously (21). Cells were suspended in the gel solution at a density of 75,000 cells per gel and 1.5 mL of mixture (per well) was poured into 12-well plates. The mixture was allowed to congeal for 30 min at 37°C, and 1.5 mL of appropriate media was added to each well (CD-FBS media containing E2 +/− calcitriol for T47D, MCF10A media +/− calcitriol). Gels were detached as previously described (22). Cultures were maintained either for 1 week to measure total cell yield or 2 weeks for morphological assessments. At each endpoint, gels were harvested and processed as described by Speroni et al (21). Briefly, to count cell numbers, cells were extracted by digesting 3D gels with collagenase and then lysed to obtain nuclei that were then counted using a Coulter Z1 particle counter (Beckman Coulter, CA). For morphological assessments, gels were harvested at 2 weeks, fixed with 10% phosphate-buffered formalin, and either embedded in paraffin for histological analyses or whole mounted and stained with carmine alum to visualize epithelial structures.

### Whole mount Analysis

Whole-mounted gels stained with carmine alum were imaged using a Zeiss LSM 800 confocal microscope for automated morphometric analysis as described by Paulose et al (23). Briefly, ^~^1 mm^2^ area of the gel periphery was imaged to a depth of ^~^100 μm. Resulting images were stitched together and analyzed using Software for Automated Analysis (SAMA) (23) and statistical analyses of morphometric parameters was performed using GraphPad Prism software.

### Picrosirius staining

Formalin-fixed, paraffin-embedded gels were sectioned using a microtome at 5μm thickness. Gel sections were then rehydrated and stained with picrosirius red solution to visualize collagen fibers and counter-stained with Weigert’s hematoxylin as described by Junqueira et al (24). Stained sections were observed under polarized light using a Zeiss Axioskop 2 Plus microscope.

### Real-Time PCR

Gels were digested using collagenase as described above, and RNA from cells was harvested using Qiagen RNeasy mini kit according to manufacturer’s instructions. Gene transcripts were quantified via RT-PCR using the Luna Universal One-Step RT-qPCR kit (New England Biolabs, MA) in an iQ5 thermocycler (Bio-rad, CA). Transcript levels were normalized to *RPL19* transcripts levels. Primer sequences for genes analyzed are as follows - *PRGAB* 5′-GAGGATAGCTCTGAGTCCGAGGA-3′ (forward), 5′-TTTGCCCTTCAGAAGCGG-3′ (reverse); *RPL19* 5′-TAGTCTGGCTTCAGCTTCCTC-3′ (forward), 5′- TCTGCAACATCCAGCTACCC-3′ (reverse); *CYP24A1* 5’-GAAAGAATTGTATGCTGCTGTCACA-3’(forward), 5’- GGGATTACGGGATAAATTGTAGAGAA-3’ (reverse).

### Statistics

GraphPad Prism and SPSS software were used for all statistical analyses. One-way ANOVA followed by Tukey’s *post hoc* test were performed to determine differences in the dose-response curves, number of elongated structures in MCF10A gels and MCF10A gel diameters; one-way ANOVA with *post hoc* Dunnett’s 2-sided *t*-test was performed for cell yields in T47D 3D gels. Kruskal-Wallis test was performed to determine differences in the morphometric parameters of structures. Chi-square analysis was performed to compare distributions of T47D epithelial structures in different volume categories. Unpaired *t*-test with Welch’s correction was used to analyze RT-PCR data. For all statistical tests, results were considered significant at *p* < 0.05.

## Results

### Effects of calcitriol on estrogen-induced cell proliferation

T47D cells exposed to different doses of vitD3 showed a decrease in cell numbers at the end of a 6-day period only when in the presence of estrogen in both 2D and 3D cultures (Fig. 1). However, the reduced cell yield was observed at the 50 nM and 100 nM doses of vitD3 alone in 2D (Fig. 1A), whereas a reduced cell yield was observed only at the 100 nM dose in 3D (Fig. 1B). Based on our inverted microscope visualization of dead “floaters” in these cultures, the decrease in cell yield, especially in 2D culture, can be attributed to cell death. In contrast, we observed an increase in cell numbers at lower calcitriol doses, such as 10 and 25 nM in 3D culture, a phenomenon that has not been previously reported. Separately, calcitriol activity in these cells was confirmed by performing RT-PCR for *CYP24A1* gene transcripts; there was a dose-dependent increase following calcitriol treatment (Supp. Fig. 1).

**Fig. 1.**
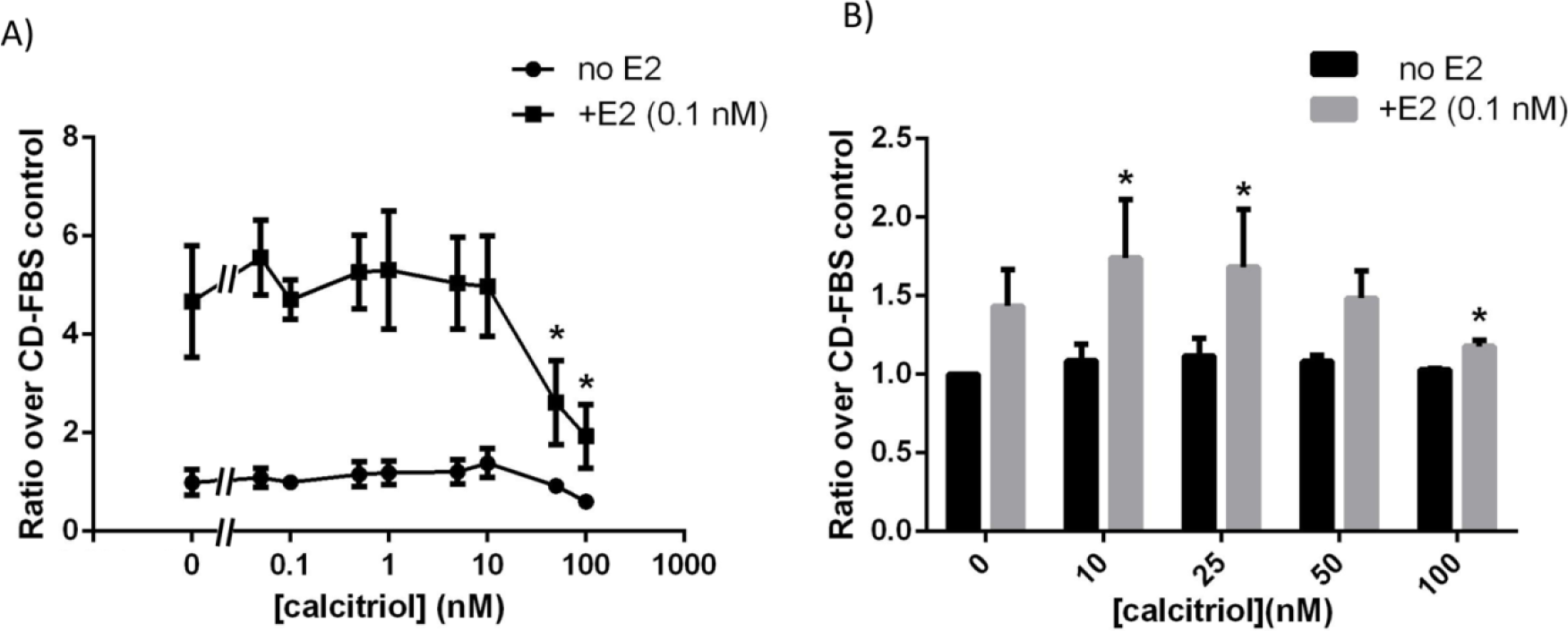
Calcitriol affected total cell yield of T47D cells differently in 2D and 3D culture conditions, and only in the presence of E2 (0.1 nM). (A) Calcitriol reduced total cell yield starting at 50 nM and higher doses in 2D culture (\**p*<0.05, one-way ANOVA). (B) Calcitriol resulted in increased cell numbers at 10 and 25 nM doses, but decreased total cell number at 100 nM dose (\**p*<0.05 compared to E2, one-way ANOVA).

### Effects of calcitriol on Estrogen-induced epithelial organization

Previous work from our lab had shown that E2 induces T47D cells to form mostly elongated shaped structures when embedded in a 3D rat tail collagen type I matrix (21). Using this same model, we investigated how vitD3 affects the organization of T47D cells in the presence of 0.1 nM E2. As shown in Fig. 2, calcitriol affects the organization of these cells, mainly by affecting the volume of the epithelial structures, in a dose-dependent manner (Supp. Fig. 2). More specifically, the change in the organization is seen through a re-distribution of different sized structures in the population – the 50 nM calcitriol dose results in a higher number of smaller structures whereas the 10 nM dose results in a slight, non statistically significant increased number of larger structures (Fig. 2B). In contrast, calcitriol alone does not seem to affect these cells when added to the hormone-depleted CD-FBS medium, thus suggesting that either the interactions with E2 are responsible for the effects observed, or the effect of calcitriol is independent of E2 but is unobservable as the cells are not proliferating in the absence of E2.

**Fig. 2.**
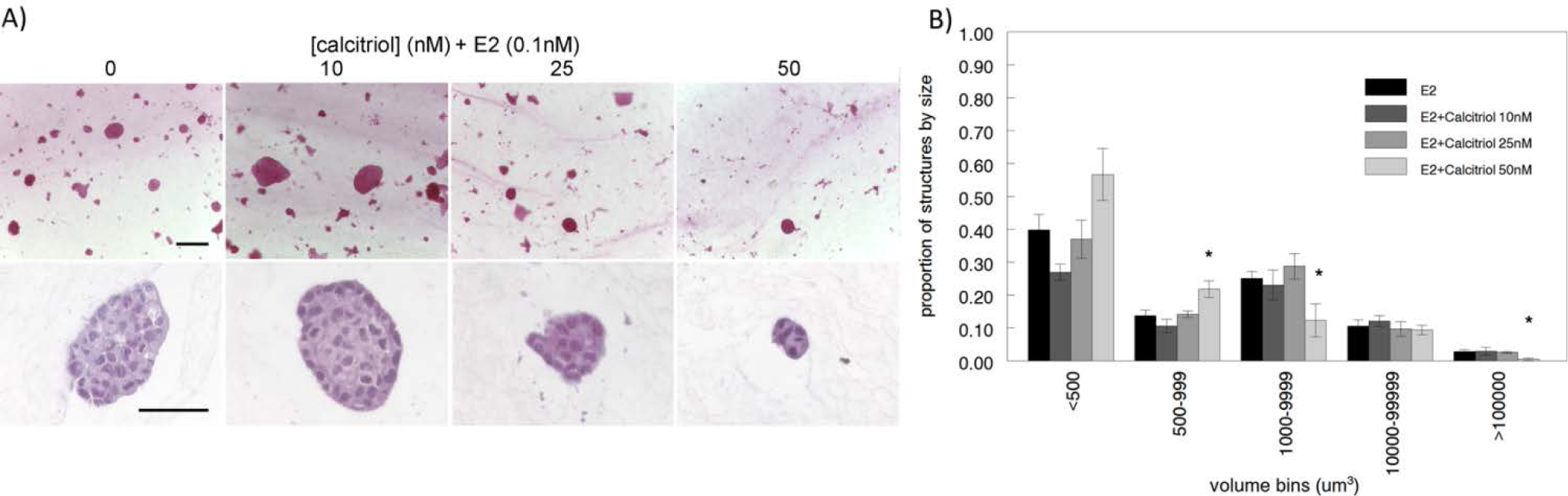
Calcitriol affected the organization of T47D cells in 3D collagen matrix in the presence of 0.1 nM E2, in a dose-dependent manner. (A) Calcitriol’s dose-dependent effects observed in carmine-alum stained whole mounted gels (top; scale bar=200 μm), and in H/E stained FFPE sections (bottom; scale bar=50 μm). (B) 50 nM dose resulted in an increased number of smaller structures with a concomitant decrease in larger structures (\**p*<0.05, chi-square).

Further investigation into vitD3’s effects on epithelial organization using our unsupervised and unbiased analysis revealed that calcitriol affects morphological parameters of the T47D epithelial structures, also in a dose-dependent manner. The 50 nM calcitriol dose resulted in a reduction in the major radius of the ellipsoid (ell_majrad; Fig. 3A) and an increase in the elongation ratio (elon1; Fig. 3B); this observation suggests that this dose results in smaller and thinner structures compared to those induced by 0.1 nM E2. In contrast, the 10 nM calcitriol dose resulted in a decrease in sphericity (Fig. 3C) and an increase in the flatness ratio (elon2; Fig. 3D); in this condition, the T47D structures are more elongated and wider compared to those in the E2 control.

**Fig. 3.**
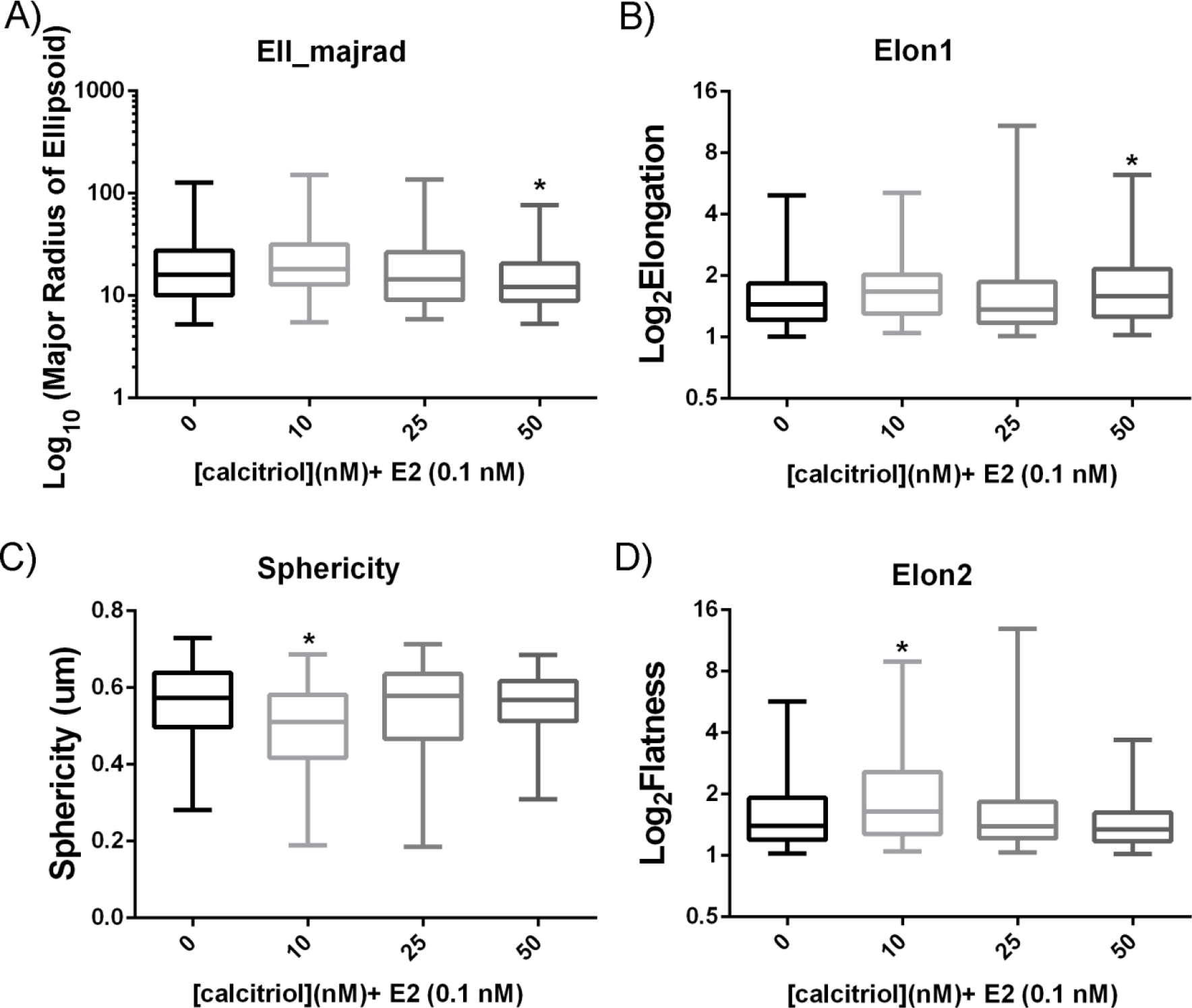
Calcitriol affected physiological parameters of T47D epithelial structures in a dose-dependent manner. Compared to 0.1 nM E2 controls, 50 nM calcitriol resulted in smaller and thinner structures as shown by a decrease in the major radius of the ellipsoid (A) and an increase in the elongation ratio (B), whereas 10 nM calcitriol resulted in more elongated and wider structures as shown by a decrease in sphericity (C) and an increase in flatness ratio (D; \**p*<0.05, Kruskal-Wallis Test.)

### Calcitriol interferes with estrogen’s transcriptional effects

Previous studies have indicated that vitD3 interferes with E2-induced gene expression in cultured cells and human breast tissue (25). To delineate whether vitD3 was interacting with E2’s activity at the transcriptional level, we performed RT-PCR to examine levels of Progesterone receptor (*PRGAB*) transcripts, which are a reliable indicator of E2’s activity both *in vitro* and *in vivo*. We observed that both 50 and 100 nM calcitriol diminished PRGAB induction by E2 by ^~^60% in T47D cells in 3D culture, as shown in Fig. 4. This indicates that calcitriol interferes with E2’s activity at the transcriptional level in our model.

**Fig. 4.**
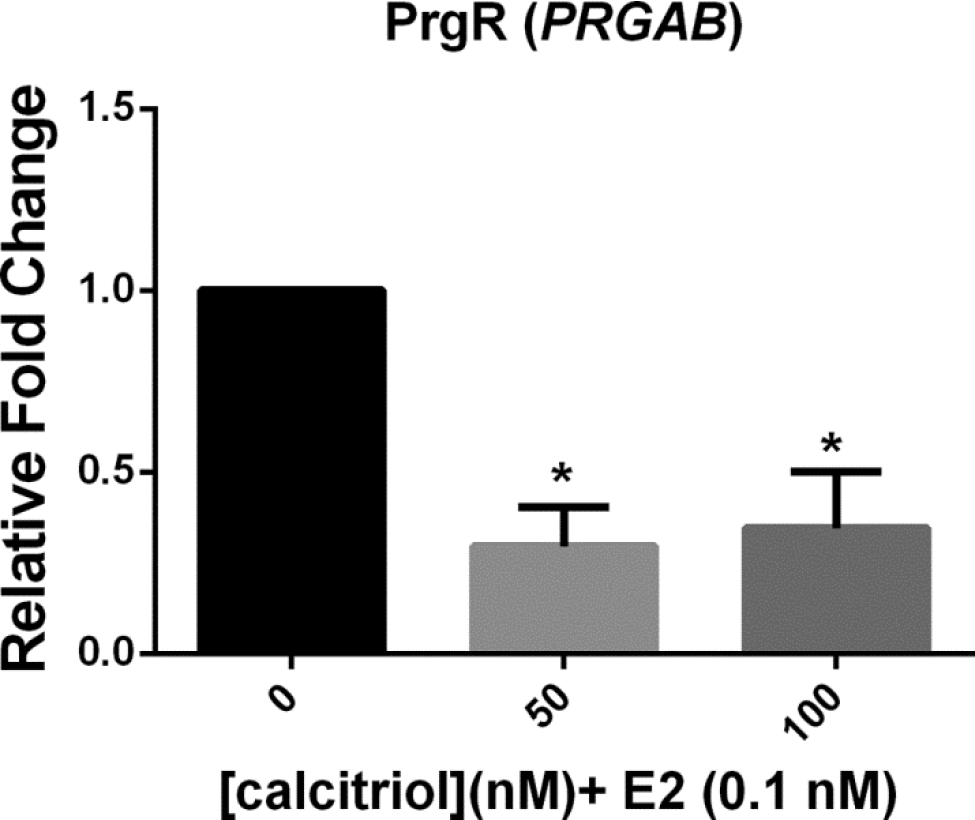
Calcitriol decreased E2-induced upregulation of Progesterone receptor expression (PrgR, *PRGAB*) at both 50 and 100 nM doses (\**p*<0.05, *t*-test). Transcripts were measured in T47D cells from 3D collagen gels after 72 hours incubation.

### Estrogen-independent effects of vitD3

MCF10A cells are considered normal breast epithelial cells because they do not form tumors when inoculated into nude/SCID mice (26) and form normal acinar and ductal structures in 3D cultures (27). To investigate whether vitD3 has autonomous effects on mammary epithelial cell proliferation and organization, we chose to investigate calcitriol’s effects on MCF10A cells under those conditions. MCF10A cells are ER negative; as shown by the induction of *CYP24A1* gene transcripts, they respond to calcitriol in a dose-dependent manner (Supp. Fig. 3). Because calcitriol exposure resulted in lower cell numbers starting at 10 nM in 3D compared to 100 nM in 2D culture, these cells appear to be more sensitive to calcitriol in 3D culture conditions (Fig. 5). MCF10A cells exhibited a monotonic dose response relationship in both cases.

**Fig. 5.**
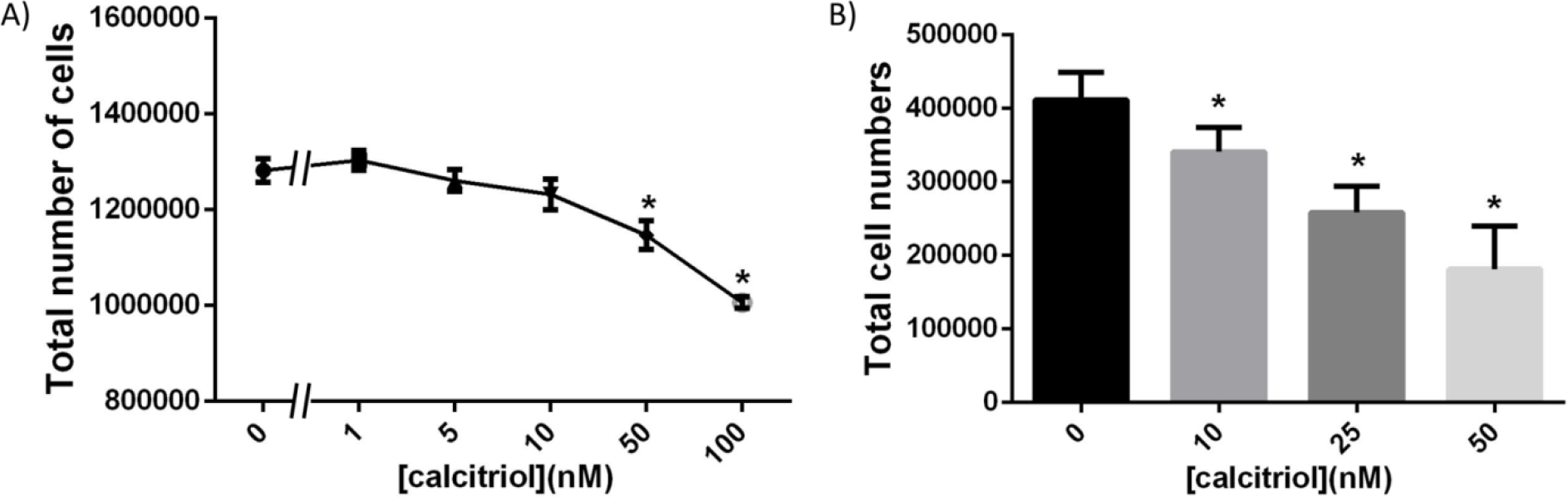
MCF10A cells showed differential sensitivity to calcitriol depending on culture conditions. (A) 100 nM calcitriol significantly decreased total cell numbers in 2D culture, but in 3D culture (B), the effects were observed starting at 10 nM (\**p*<0.05, one-way ANOVA).

### Autonomous effects of calcitriol on epithelial organization

MCF10A cells embedded in bovine type I collagen formed predominantly ductal structures with few acinar structures after 2 weeks in culture (27). Using this model, we investigated vitD3’s effects on the organization of these cells; however, we utilized a rat tail type I collagen matrix (Fig. 6). Previous work from our lab (28) has shown that epithelial organization in 3D culture is influenced by the species of the collagen type I used in the extracellular matrix formulation. Consistent with those and other findings (29), we observed mostly acinar structures with few ductal structures in a rat tail type I collagen matrix (Fig. 6A). Interestingly, we also observed that calcitriol increases the number of elongated structures in 3D culture, with the highest number observed at the 10 nM dose, followed by a reduction at higher doses (25 & 50 nM, Fig. 6B).

**Fig. 6.**
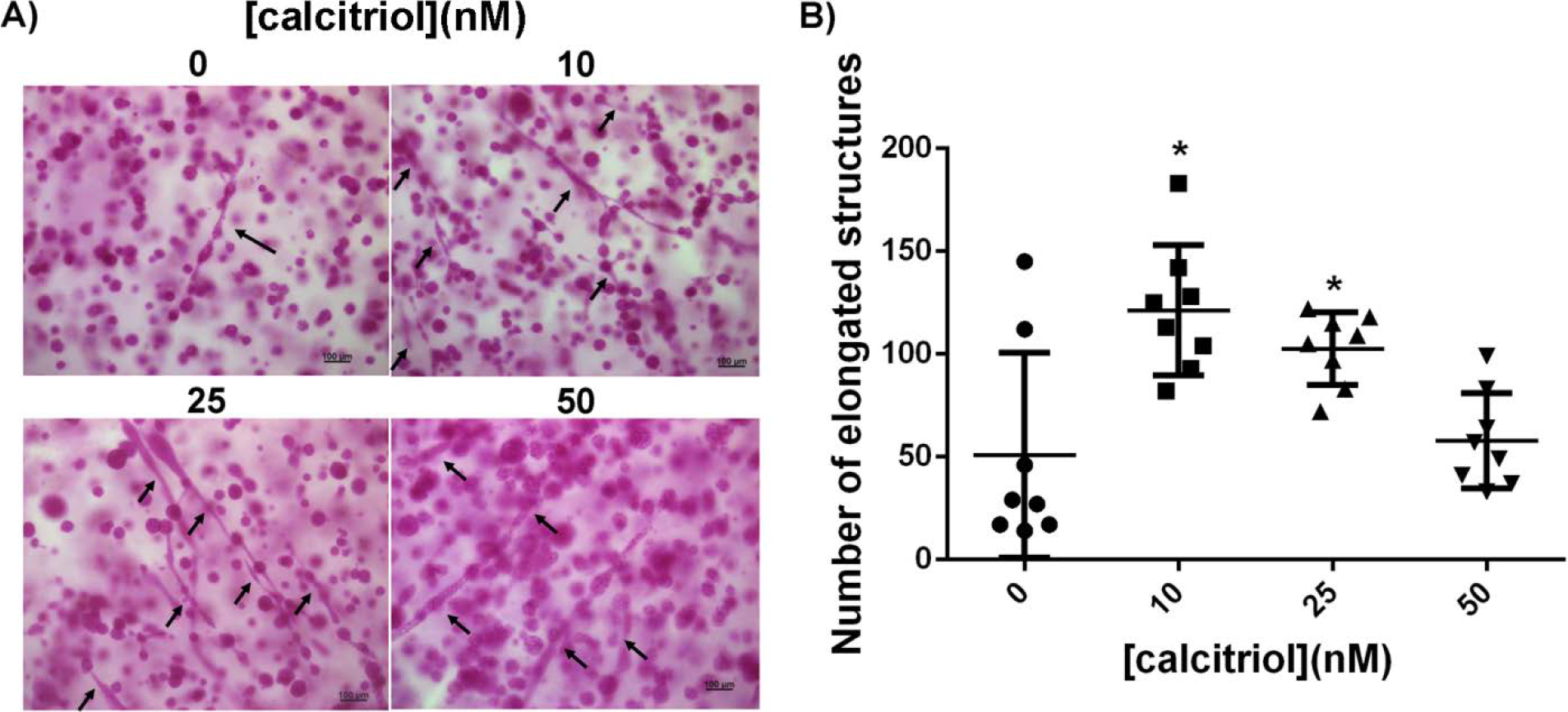
Calcitriol treatment resulted in increased number of elongated structures in a non-linear fashion. (A) Carmine-alum stained whole mounts showing elongated structures (arrows; scale bar= 100 μm). (B) Quantification of elongated structures in gels show that 10 nM calcitriol resulted in the highest number with consequent decline in higher calcitriol doses (\**p*<0.05, one-way ANOVA).

### Effects of calcitriol on MCF10A epithelial organization

Confocal images of MCF10A 3D gels were analyzed for changes in the morphological parameters of the epithelial structures upon calcitriol treatment. Exposure to calcitriol resulted in flatter and less spherical structures; these effects were significant at the 50 nM dose (Fig. 7). Calcitriol also resulted in a decrease in the volume of these epithelial structures, most significantly at 50 nM dose (Supp. Fig. 4), which can be attributed to the increase in flatness with increasing calcitriol doses.

**Fig. 7.**
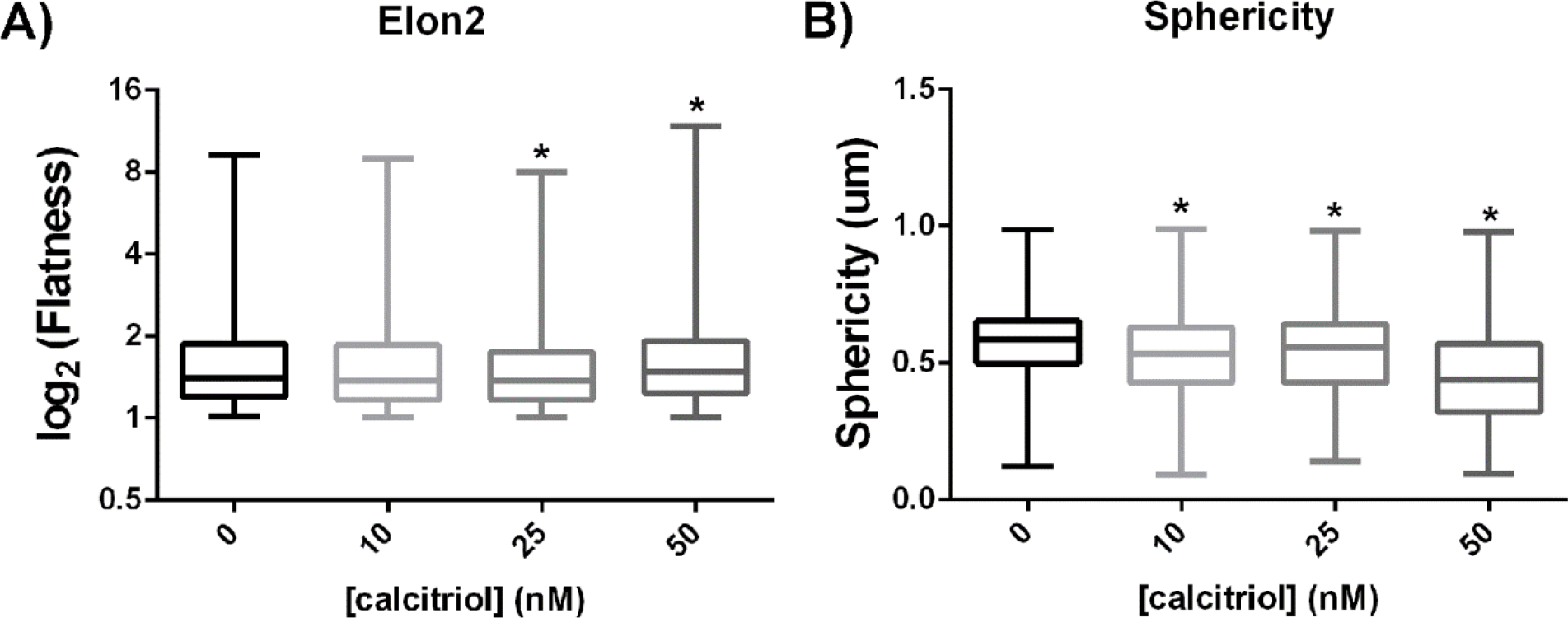
Calcitriol treatment resulted in (A) flatter and (B) less spherical MCF10A epithelial structures in a type I collagen 3D matrix when cultured for 2 weeks (\**p*<0.05, Kruskal-Wallis.)

### Effects of calcitriol on collagen organization

MCF10A cells organize collagen fibers in the 3D gels, as a prerequisite for organizing into ducts or acini (30). Because of the changes in the morphological parameters of the epithelial structures, we investigated collagen fiber organization in 3D gels using picrosirius staining. Polarized light microscopy of FFPE sections stained with picrosirius revealed that calcitriol has a non-monotonic effect on collagen fiber organization in this 3D model (Fig. 8). Calcitriol at 10 nM dose reduced the number of organized collagen fibers. In contrast, 25 and especially 50 nM doses showed an increase in the amount of organized collagen fibers. Additionally, at these doses, organized fibers were more uniformly distributed throughout the 3D gels, especially in areas distal from epithelial structures (data not shown). These observations may explain why calcitriol treatment also resulted in increased contraction of the 3D gels in a dose-dependent manner after 2 weeks in culture (Supp. Fig. 5).

**Fig. 8.**
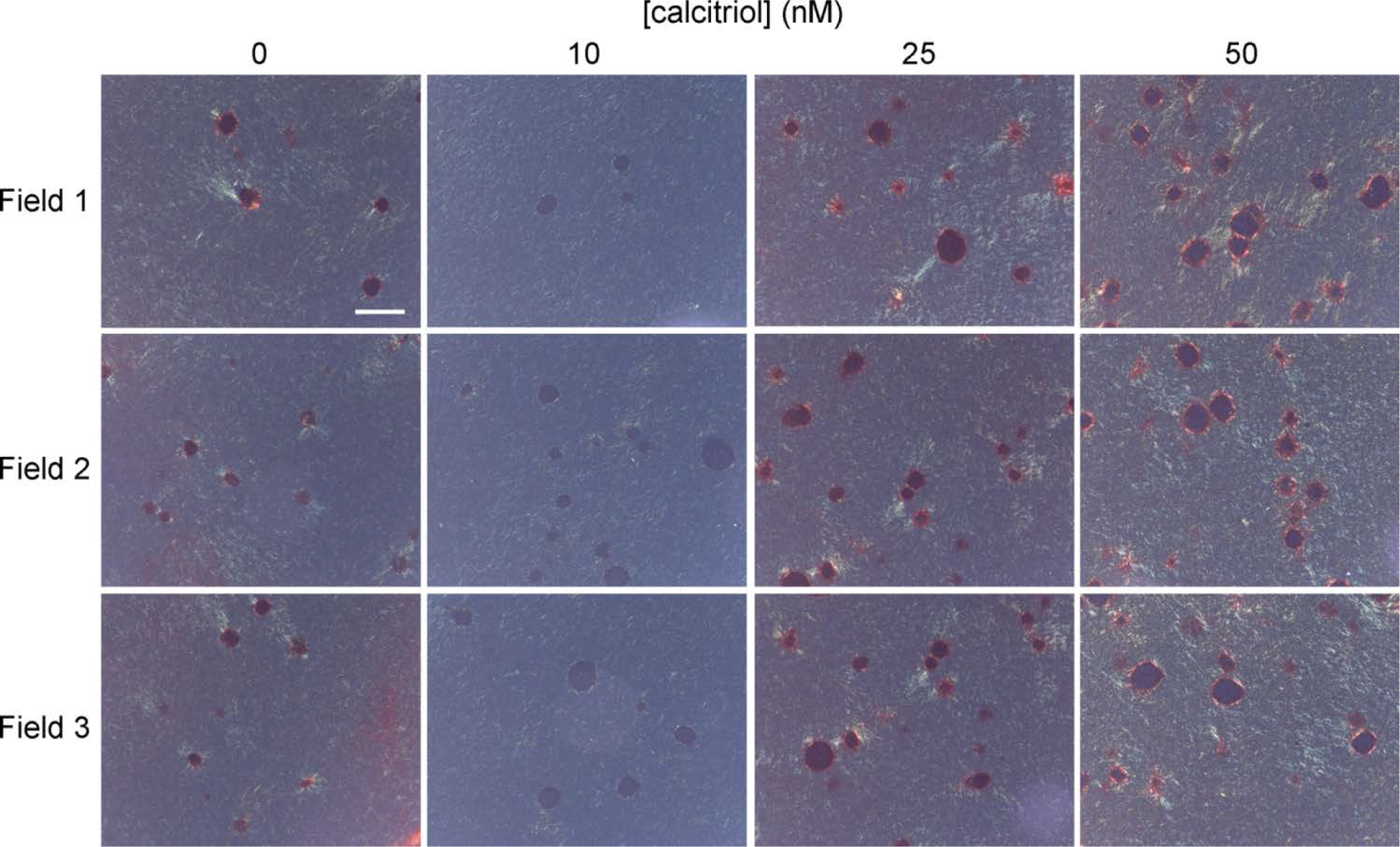
Calcitriol exposure resulted in a different collagen organization in MCF10A 3D gels. See representative images from three different fields of view for each treatment group (scale bar=100 μm)

## Discussion

Despite evidence linking vitD3 deficiency to increased risk of breast cancer and worse clinical outcomes in patients, randomized clinical trials have yet to confirm the efficacy of vitD3 as a preventive or therapeutic option in this disease (8). Experimentally, while VDR KO mice do not develop tumors spontaneously, mammary glands from these mice exhibit a striking phenotype of excessive and precocious development at key stages (15,16). This suggests that vitD3 plays an important role in the development of the normal mammary gland. When considering that carcinogenesis is “development gone awry” (7), an understanding of the role of vitD3 in this process may provide worthy therapeutic options for breast cancer patients.

The VDR is expressed in the mammary gland at the different stages of postnatal development that are largely influenced by the mammotropic hormones E2, Prg and Prl. These hormones have well characterized effects on the morphogenesis of the gland; for example, E2 stimulates ductal elongation, Prg increases lateral branching and Prl induces alveologenesis (17). Although the VDR KO phenotype of the gland has been described (15,16), no reference has been made so far to the interactions between E2, Prg and Prl, and vitD3 in a 3D environment in which morphogenesis takes place. To fulfill this need, we have utilized two different 3D culture models to tease out vitD3’s effects that are either dependent or independent of its interactions with E2. We noticed that calcitriol exhibits a non-monotonic dose response only in 3D cultures, a phenomenon not previously described. This is in line with current knowledge regarding steroid hormone activity and also favors the notion that vitD3 functions as a steroid hormone.

In the presence of estrogen in 3D culture, calcitriol increases total cell yield at 10 nM dose whereas it decreases total cell yield at 100 nM dose (Fig. 1B). This reduction in cell yield can be attributed to cell death given our observation of floater cells in both 2D and 3D cultures. Comparable evidence was found when the role of vitD3 was explored in apoptosis (8,25). We also observed that calcitriol constrains the effects of E2 on mammary epithelial morphogenesis without affecting total cell yield, more specifically on the organization of epithelial ductal structures in 3D conditions. Consistent with our finding, the mammary glands of *CYP24A1* KO mice, which cannot metabolize calcitriol, exhibit stunted development (31); Zinser et al (16) report that VDR KO mammary glands exhibit increased ductal elongation at puberty. Of note, in both of these models, the proliferative capacity of the epithelial cells was not affected. Given that E2 is responsible for ductal elongation during puberty, our results recapitulate vitD3’s activity *in vivo* and confirm that calcitriol constrains the effects of estrogen on ductal elongation without affecting cell death or proliferation.

During puberty, the mouse mammary gland epithelial ducts elongate under the influence of E2. However, only 15-20% of epithelial cells in the gland are ER-positive at that time (32), with this protein being expressed in the interior luminal layer of epithelial cells and not in the outermost cap layer. A similar pattern of VDR expression in the mammary gland epithelium at puberty has been reported, with most expression observed in the trailing edge of the terminal end buds and lesser expression in the cap cells (15). Therefore, while E2 appears to directly affect 15-20% of epithelial cells during puberty, its effects are observed at the tissue level.

In order to investigate whether vitD3 autonomously affects epithelial cells beyond its interactions with E2, we utilized an estrogen-independent 3D culture model. MCF10A cells, considered to portray a “normal-like” behavior *in vitro*, organize into mostly acinar and form some ductal structures in the 3D collagen matrix (27). We observed that vitD3 retains its non-monotonic effects on morphogenesis even in an estrogen-independent 3D culture model. We showed that this organization of MCF10A cells is affected in a dose-dependent manner when treated with calcitriol in 3D cultures. These cells showed greater sensitivity to calcitriol in 3D when compared to 2D cultures, with cell death increasing in a dose dependent manner upward from a 10 nM dose (Fig. 5).

Mechanical forces are the main mediators of shape during morphogenesis. Previously, we have shown that mammary epithelial cells embedded in a type I collagen matrix manipulate the collagen fibers around them in the process of organizing into complex shapes such as ducts and acini (21,30); these epithelial cells exert mechanical forces that act on collagen fibers and on other cells. As fibers organize, they constrain the cells on their ability to move and to proliferate (33). We have also shown that hormones distinctively influence the way epithelial cells organize collagen fibers, and consequently determine the shape of the structures formed (21,34). Based on these results, we hypothesized that vitD3 would also affect fiber organization. MCF10A cells treated with increasing concentrations of calcitriol in 3D cultures increase the contraction of gels. Treatment with calcitriol also decreased the number of cells in the gels in a dose-dependent manner. Gel contraction is dependent on the number of cells present in the gel and on the manipulation of collagen fibers by the cells. While the lower number of cells can account for the smaller sizes of structures observed in the 3D gels treated with calcitriol, this does not explain the increased contraction of these gels. Picrosirius staining revealed that even though there are fewer cells in the gels at 50 nM dose, there is a more uniform distribution of organized fibers throughout the gel (Fig. 8). As our lab and others (35) have described, organized fibers are responsible for transmission of forces and more organization of fibers leads to increased anisotropy in the 3D environment. The increased contraction of the gels can therefore be explained by the transmission of forces across long distances by the cells, a phenomenon previously reported (36).

We also observed that at 10 nM dose of calcitriol there was the least amount of organized fibers and the greatest number of elongated structures (Fig. 8). On closer observation, the calcitriol treated gels contained a lower number of branched, elongated structures (ductal, tubular) and a higher number of unbranched, elongated structures (cord-like). Additionally, when compared to untreated gels, increase in calcitriol dose resulted in shorter and thinner elongated structures. We have previously reported that MCF10A cells embedded in a collagen type I matrix form ductal/tubular structures in the periphery of the gel and cord-like structures in inner areas (22). In the calcitriol treated gels, in addition to the inner areas, cord-like structures were also observed in the periphery. The arrangement of collagen fibers by the cells is affected by a multitude of factors that include physical constraints. Therefore, in order to fully elucidate the differential organization of structures depending on the calcitriol dose used, additional measurements of local biomechanical parameters in the calcitriol-treated 3D gels is required.

All models, by definition, are simplified versions of the object being modeled. They are used precisely because they reduce the number of variables considered relevant to explain a phenotype. Thus, like all experimental models, 3D culture models have their limitations. For example, the use of established cell lines, which are considered rather stable is dictated by the limitations of using freshly isolated primary cells. Isolated human primary cells are not efficient in forming biologically relevant structures in collagen or ECM matrices *in vitro*; only a small percentage of them express mammotropic hormone receptors and they lose their potential to form structures shortly after being placed in culture (37–39). Considering these inherent limitations, we have utilized human breast epithelial cell lines such as T47D and MCF10A to create more robust, consistent and complementary models that would still mimic the mammary gland morphogenesis observed *in vivo*.

Future work should incorporate findings from *in vitro* 3D models and test them in an *in vivo* model. To that end, findings that calcitriol constrains the action of estrogen can be incorporated into a mammary gland transplant model between VDR KO and wild type animals to investigate the role of the stromal and epithelial compartments in mediating vitD3’s effects during pubertal development. Similarly, a fetal mammary gland *ex vivo* culture model (40) can also be utilized to more comprehensively understand the role of vitD3 in early development.

Here, we have shown that calcitriol, at physiologically relevant doses, have effects beyond cell death and proliferation described in the current literature. This study highlights the role of vitD3 as a morphogen to the extent that calcitriol contributes to the proper shape formation of the mammary gland development. Disorganization of the tissue architecture during early developmental phases has been shown to contribute to tumor formation in the mammary gland in adult life (6,41). This study provided a more detailed understanding of vitD3’s role in normal mouse mammary gland development and a lead on how vitD3 deficiency might contribute to increased breast cancer risk.

## Acknowledgements

We would like to thank Cheryl Schaeberle for her help with statistical analyses and a critical reading of this manuscript.

